# IRIS: an accurate and efficient barcode calling tool for *in situ* sequencing

**DOI:** 10.1101/2020.04.13.038901

**Authors:** Yang Zhou, Hao Yu, Qiye Li, Rongqin Ke, Guojie Zhang

## Abstract

**Summary:** The emerging *in situ* RNA sequencing technologies which can capture and amplify RNA within the original tissues provides efficient solution for producing spatial expression map from dozens to thousands of genes. Most of *in situ* RNA-seq strategies developed recently infer the expression patterns based on the fluorescence signals from the images taken during sequencing. However, an automate and convenient tool for decoding signals from image information is still absent. Here we present an easy-to-use software named IRIS to efficiently decode image signals from *in situ* sequencing into nucleotide sequences. This software can record the quality score and the spatial information of the sequencing signals. We also develop an interactive R shiny app named DAIBC for data visualization. IRIS is designed in modules so that it could be easily extended and compatible to new technologies.

**Availability and implementation:** IRIS and DAIBC are freely available under BSD 3-Clause License at: https://github.com/th00516/ISS_pyIRIS.

## Introduction

Spatial transcriptomics is an emerging field that aims to characterize the gene expression profiling together with the spatial context of the tissues(Burgess, 2019; Stark, et al., 2019). It offers solutions to address many fundamental questions on cellular function. Several *in situ* RNA-seq technologies have been developed recently allow the high throughput detection of gene expression *in situ* with the high resolution fluorescence image (Chen, et al., 2015; Ke, et al., 2013). These technologies usually involve the visualization and quantitative analyses of transcriptome with spatial resolution from the fluorescence images of tissue sections. However, there is no any software to decode the sequencing signals from images, which limits the application of these new technologies for downstream analyses. Here, we demonstrate an open source software IRIS (Information Recoding of *In situ* Sequencing) to decode image signals into nucleotide sequences along with quality and location information. We also present an R shiny app DAIBC (Data Analysis after ISS Base Calling) for interactive visualization of called results. IRIS shows good performance in both data processing efficiency and accuracy at gene expression and location levels. We also designed it in modules so its compatibility could also be further extended to other technologies.

## Implementation

We employ image and directory structure of *in situ* sequencing (ISS) (Ke, et al., 2013) as our default input data structure. Images are organized as split channels and sorted in different cycles. Each cycle includes five image channels, which are marked by the fluorescent dyes, Y5, FAM, TXR, Y3, DAPI, representing dyes for base A, T, C, G and nucleus, respectively (Fig. 1A). Different with images in traditional next generation sequencing (NGS), ISS images contain not only fluorescent spots, but also background like nucleus and cytoskeleton (Supplementary Fig. 1), which produce background noise that need to be filtered before decoding. Thus, we took several steps including registration, blob detection and connection to decode image signals into barcodes.

**Figure 1.**
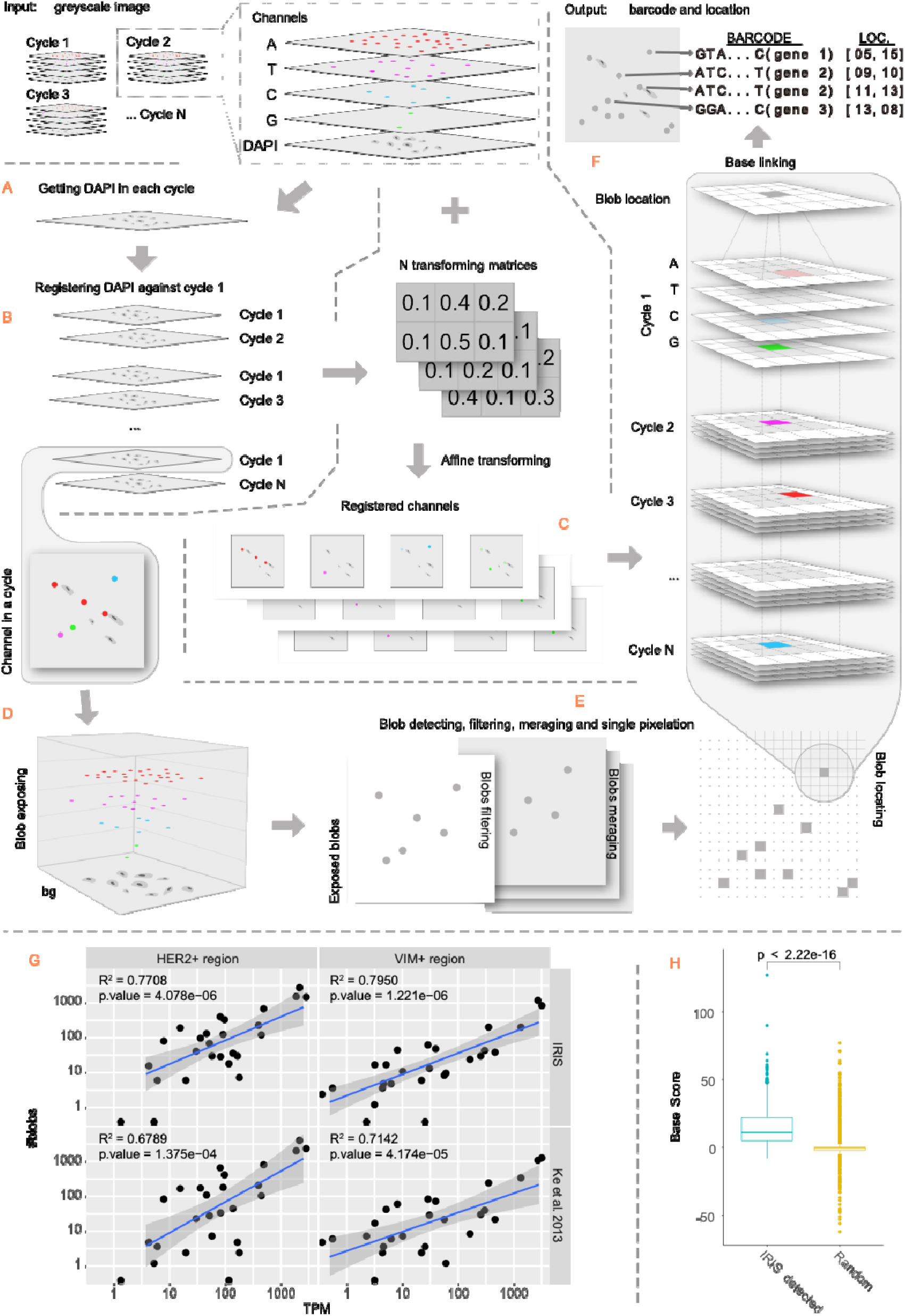
General workflow and evaluation of IRIS. We import all images from all cycles as matrices and store them into a stack data structure. (A) DAPI images from different cycles are used as the representative image of each cycle for registration. (B) DAPI of each cycle is registered with DAPI of cycle 1 to obtain the transform matrix for each cycle. (C) These transform matrices are used to register all channels in their own cycles. (D) Blobs in each registered channel in all cycles are exposed by tophat transformation, and their coordinates are recorded. (E) Blobs’ coordinates from all cycles are map into a reference layer for redundancy removal and to generate a coordinate reference of all blobs. (F) This reference is used to connect all bases called from registered channels of different cycles. At last, base calling information is produced as output, composing of five columns for each blob, including blob ID, barcode sequence, quality, row and column in cycle 1 DAPI image. (G) The correlation between the expression signal detected by IRIS and TPM inferred from RNA-seq in HER2+ and VIM+ region. (H) The base score distribution of blob detected by IRIS is substantially higher than the score from random pixels.

### Intermediate data structure and images registration among different cycles

Because the positions of cells and transcript amplification products in different cycles can be shifted during experiment operation, the first step of IRIS is image registration, which aligns images from different cycles to the same coordinate system. Images from the same cycles doesn’t need to be registered as their differences are mainly raised from exposure time. During registration, key points are first identified from the images and used as makers to align images from different cycles. Then transformation matrices are calculated based on the matched key points pairs between two images and used to align images from different cycles to the same coordinate system.

In order to reduce error in registration, we first remove noise in each image. A low-pass filter is performed to filter out pixels with the 40% highest signal frequency after Fast Fourier Transformation. We by default implement of ‘ORB’ algorithm (Rublee, et al., 2011) to collect the coordinates and measure the scales and orientations of key points (i.e. description of key points). We further identify matched key point pairs with similar description between every image from cycle N to image from cycle 1 with k-Nearest Neighbor (kNN) on the description matrix of key points (Altman, 1992) (Supplementary Fig. 2). Then we iteratively filter out matched key point pairs outlier with large distance that might be false-positive caused during matching process. The final key point pairs are used to calculate homography after being sorted by pair distance. This process generates one transform matrix for each cycle, which can be used to register every channel in each cycle respectively (Fig. 1C, Supplementary Fig. 2). In some ISS technologies, DAPI is used to capture nucleus structure thus is present in all cycles. In each cycle it harbors more key points thus provides an ideal information for image registration. If this image is available, we make it as the default images for registration in IRIS (Fig. 1A).

### Blobs detection in each cycle

Hybridization signals are presented as light-spot of certain size under dark background in the image, thus can be treated as blobs in computer vision area. Blobs of registered image in each channel will be exposed by tophat transformation under 15×15 ellipse kernel. We roughly detected blobs from each exposed image with ‘SimpleBlobDetector’ of OpenCV (Bradski and Kaehler, 2000). To obtain a non-redundant blob set for each cycle, the detected blobs from all channels in the same cycle will be mapped to a single size-equivalence layer with no background to obtain a non-redundant blob set for each cycle (Fig. 1D and E, Supplementary Fig. 3)

A crucial feature of a real blob is that pixel grayscale increases dramatically in its core region compared with its periphery. While previous detection step could expose a number of blobs, it would also include some false-positive because some regions in the images might have elevated background brightness or noise surrounding which might be overexposed and falsely detected as blobs. To reduce false-positive, for each blob, we utilize the difference between the mean of grayscale in the core region (4×4) and that in the periphery (10×10), which reflects the signal strength difference between candidate blob and its surrounding background, defined here as ‘base score’. Subsequently, for each blob, base scores from different channels in the same cycle will be sorted, and the base channel with the highest base score is considered as the true base of this cycle. We further calculate the error rate P as 1-q, where q is defined as the maximum base score (i.e. the score of the assigned base in the cycle) divided by the sum of score of all channels produced in that cycle. Then we calculate the base calling quality Q by *Q* = −10 log_10_ (*P*) similar as the Phred quality score in NGS platform.

### Bases sequence connection among different cycles

Linking bases at the same location from different cycles to generate barcode sequences is a crucial and the most time-consuming step. Blobs from different cycles might not be completely overlap with each other, thus we collect all detected blobs from all cycles and project them into a new layer called ‘reference layer’ and detect blobs on this layer again to remove redundancy. This reference layer should cover all potential blobs without redundancy. Then, we take each blob in reference layer and connect bases from the first cycle to the last at each blob location. When there’s no blob detected at the location in one cycle, we add an ‘N’ with quality of one. In addition, although registration at the first step aligns most regions of images, blobs’ location might not be accurate at pixel level. To resolve this problem, we first project the location of each blob in reference layer to cycle N (defined as the searching center), and search for any candidate linked base in a 6×6 region near the center (Fig. 1F). Error rate for each candidate base detected from the searching process would need to be calibrated. The distance from a blob center at reference layer to the pixel of searching center at cycle N is defined to be one, and the distance from a searched pixel at cycle N to the searching center is defined to be D, then, we could adjust the error rate for the base at cycle N by multiplying the raw rate by 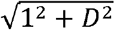 (Supplementary Fig. 4). Thus, the longer distance between the candidate and the searching center is, the harder for the error rate will be penalized. The candidate with the smallest penalized error rate thus is selected as the base of the position in cycle N. All called sequences are included in the final raw output even when there’s one or more ‘N’s. And users could further filter the sequences based on the designed barcode list, base quality, etc.

## Application and evaluation

We utilized the published ISS data (Supplementary Table 1) to evaluate the performance of IRIS. When dealing with the co-culture of human and mouse cells sample (HM), IRIS could detect 225 barcodes with 88.58% true positive rate (TPR). Specifically, when dealing with blob-dense regions, IRIS could detect all blobs without merging the spatially close ones (Supplementary Fig. 5). TPR could reach 72.42% in another breast tumor slice sample, where IRIS also achieved a higher correlation with RNA-seq expression level than previous result (Fig. 1G). In both cases, base score distribution of IRIS detected blobs was significantly higher than that of random pixels (Fig. 1H, Supplementary Fig. 6), implying the high accuracy rate of IRIS’ detecting blobs. Moreover, IRIS could deal with ISS data efficiently. For example, when dealing with the HM dataset, including a total of 20 images each with 1330×980 resolution, IRIS could finish the run (4 cycles) in approximately 11.7 CPU seconds in a one-line command. And when dealing with larger dataset like the breast tumor slices, which included 80 images each with c.a. 1390×1040 resolution, it took approximately 87.5 CPU seconds in a parallel and a total of 725.2 CPU seconds for all 16 slices (Supplementary Table 3). We also found the computation performance was affected more by the number of detected blobs rather than the total input image size (Supplementary Fig. 8).

IRIS can also handle image data generated by other ISS technologies by adding the corresponding input parser modules. For example, MERFISH utilizes binary barcodes to represent genes, so two instead of four channels are treated in each cycle (Chen, et al., 2015). After minor modification of the input data structures, the following steps can be unified and barcode sequences and locations could be called automatedly.

## Contribution

G. Z. designed and administered this project; Y. Z., H. Y., Q.L., G.Z wrote this article; Y. Z. took part in the development of IRIS, developed the code of DAIBC and performed evaluation; H. Y. designed and developed IRIS, and took part in the development of DAIBC and evaluation; R. K. provided ISS data set and suggestions for the development and evaluation of IRIS and DAIBC; G. Z. provide all the funding and resources for this work.

## Acknowledgement

We thank for Linsen Li for suggestion in registration and Shaohong Feng for manuscript revision.

