## Supplementary Information for "IRIS: an accurate and efficient barcode calling tool for *in situ* sequencing"

#### Supplementary method

As *in situ* sequencing technologies captured gene expression as well as location, we evaluated IRIS' calling accuracy in both aspects. Accuracy of gene expression level was evaluated in two ways: 1. called barcode true positive rate (TPR), defined as the expected called barcodes divided by the total number of called barcodes; and 2. correlation coefficient between blob number of the called genes and their expression level inferred by RNA-seq. To assess barcode spatial accuracy, we first performed sampling from IRIS detected barcodes and random pixels of cycle 1 merged image (without DAPI). Base score of each barcode or random pixel was calculated by subtracting the mean grayscale in a 4x4 core region to the mean grayscale in a 10x10 region. Because sequencing signal is featured as fluorescence, we would expect that the base scores of called barcode set with spatially accuracy are significantly higher than that of random blobs (Supplementary Fig. 6). One-sided t-test was performed to test the difference between the two base scores set.

For HM data, we compared the results of Ke's pipeline run with CellProfiler (v2.1.1) (Kamentsky, et al., 2011) against that of IRIS with default parameters. Due to the independent registration in the two processes, we manually curated the detected barcodes' location in Ke's pipeline by moving them 12 pixels left and 3 pixels down and used the same cycle 1 merged image as background in plot for comparison. In addition, we evaluated the spatial accuracy by comparing base scores between IRIS detected barcodes and random blobs.

For breast tumor slice data, we used expression level inferred from RNA-seq data as standard to evaluate the quantification accuracy of IRIS and Ke's pipeline. RNA-seq reads of normal breast cell lines (SRR015270-SRR015273) and breast tumor cell line (SRR015342-SRR015348) were mapped to human genome (Ensembl release 87) by HISAT2 (v2.0.5) (Kim, et al., 2015) with default parameters. Only uniquely mapped reads were used in TPM calculation. Barcodes with mean base quality $\leq 10$, homopolymers or barcode containing 'N's were removed before evaluation. For each gene, we calculated its expression (unit in blob number) in every 100x100 pixel window, and normalized the expression in two steps: 1) among windows by the maximum blob number of all genes in each window, and 2) among genes by the normalized maximum blob number of each gene. Pairwise Manhattan distances between every two windows were used for K-means clustering (K=3, R package 'amap' with parameter "iter.max = 1000, nstart = 100"). We defined the clusters with the highest mean blob number of VIM and HER2 to be VIM+ and HER2+ regions, respectively (Supplementary Figure 7). Blob number of each gene in VIM+ and HER2+ regions were used to compare with expression inferred from RNA-seq of normal breast cell line and breast tumor cell line, respectively. Pearson's correlation coefficient was calculated between $log_{10}$(TPM+0.5) and $log_{10}$(#blobs+0.5). We also evaluated the spatial accuracy of breast tumor slices' called barcodes using base scores in the same way as above.

For MERFISH, we converted the raw DAX into TIF by storm_anlaysis (https://github.com/ZhuangLab/storm-analysis) and ImageJ (Rueden, et al., 2017). Binary barcodes were called by IRIS with optimized paramters. No filtration on quality or 'N' in barcodes was applied to called barcodes as they are encoded in a binary way. Only barcodes with Hamming distance to designed barcode ≤1 were kept. We also ran Zhuang's MERFISH analysis (https://github.com/ZhuangLab/MERFISH_analysis) for comparison (Supplementary Fig. 9).

### Supplementary discussion

We designed and implemented IRIS to call barcode sequences and location in high accuracy and performance. The framework for transforming ISS images into barcodes are clearly defined in modules, while each of them could be independently substituted or optimized to improve performance and compatibility to images generated by other technologies.

Moreover, we suggest gene quantification accuracy could further be improved via standard experimental procedures. Although many mature key points-based strategies have been published, all of them are dependent on detecting enough number of key points (e.g. rigid transform requires at least 4 key point pairs), which requires images for registration to be similar enough. Unfortunately, the experimental procedures of ISS are still not standardized (Ke, et al., 2013). For example, as a manual process, images from different cycles might differ in brightness due to the various exposure time, which further lead to discrepancy of contour at the same location. This registration imperfection would impair the following base connection thus reduce the sequence accuracy of detected barcode (see the case in Supplementary Fig. 10) and effect later quantification. Moreover, unstandardized procedure will lead to discordant brightness among channels within the same cycle. This will interfere the process of blob detection and may lead to error in base score and error rate calculation. We believe that expression quantification would be improved if standard experimental procedures are taken.

How to define base-calling accuracy is also crucial in location and quantification evaluation. For location evaluation, the best way is to manually check the overlapping between called barcodes location and the fluorescent signals of the raw images. However, it becomes too inefficient and subjective if image contains too many blobs. In our case, we calculated the difference of mean grayscale between core and peripheral regions, despite that it is not precise enough. Regarding to quantification, we achieved approximate 0.80 correlation coefficient with RNA-seq data, but there are still some defects in evaluation method. RNA-Seq data (e.g. TPM, FPKM, RPKM) or smFISH is often used as the golden standard to evaluate ISS quantification accuracy (Chen, et al., 2015; Ke, et al., 2013). However, compared with bulk RNA-Seq, samples of *in situ* sequencing just are thin slices or contains only a few cells, thus its expression level might not be able to be reflected from bulk cells.

### Supplementary Figures

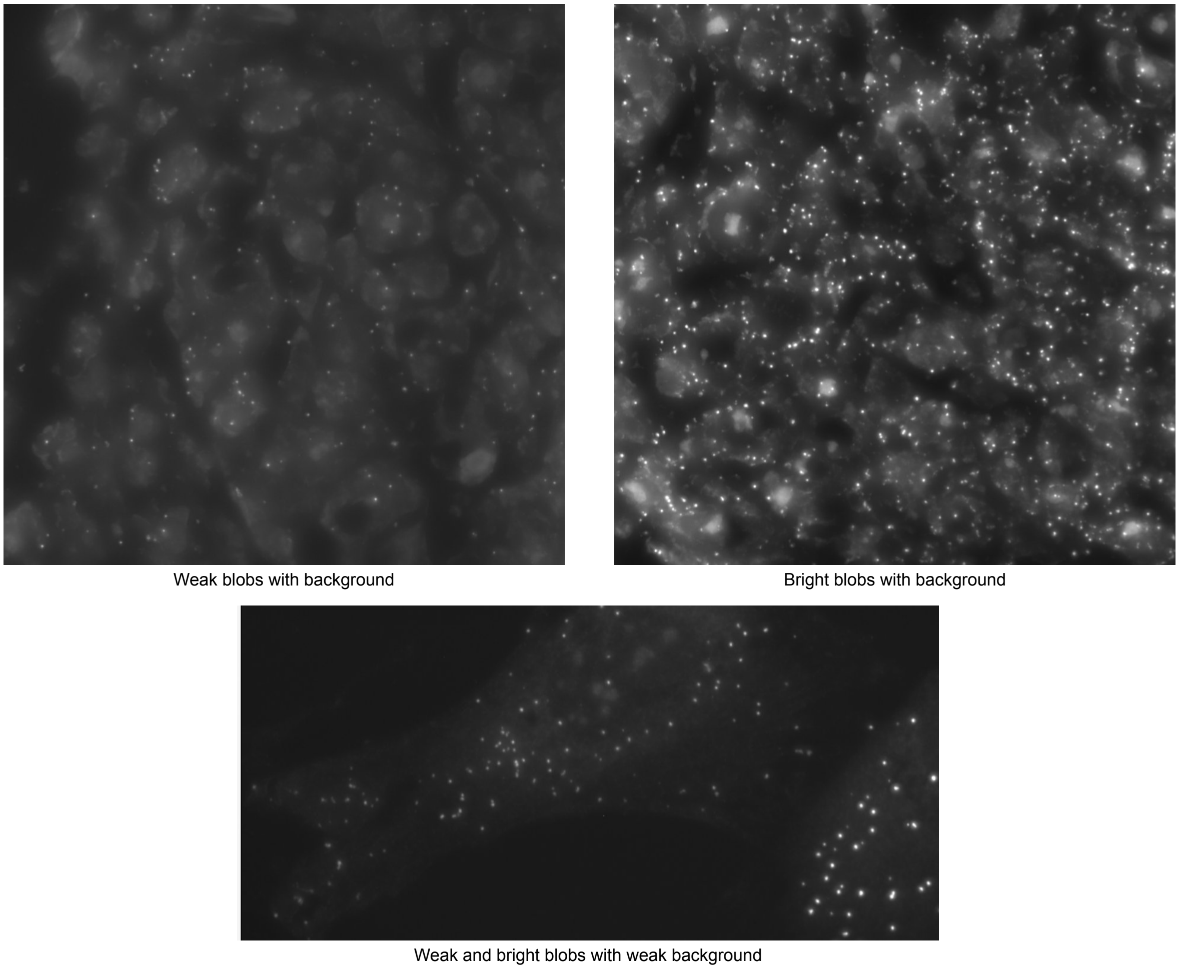


**Supplementary Figure 1. Examples of blobs with background noise.** In addition to background noise from cell structures, blobs' brightness might vary among regions, even within the same channel.

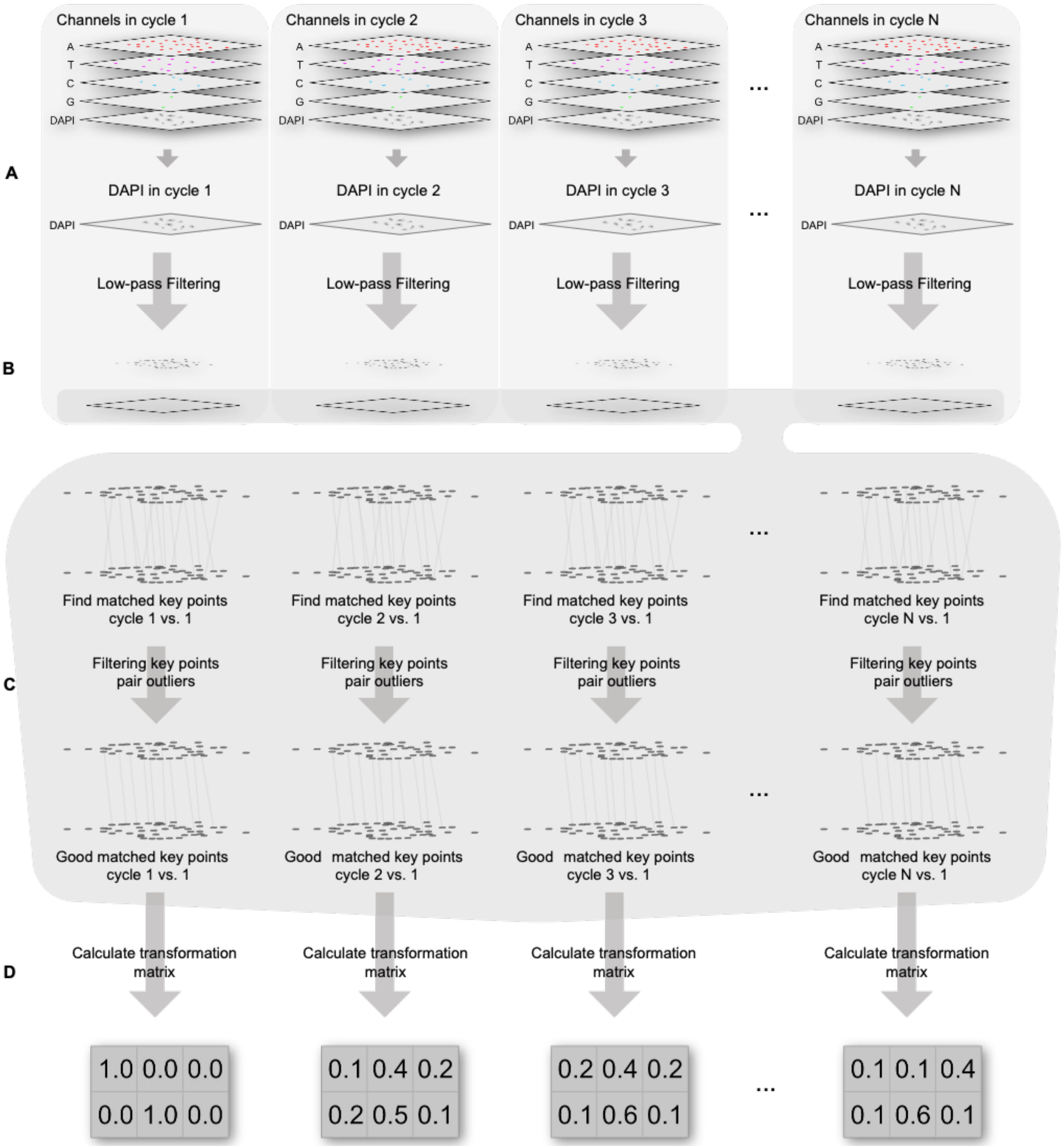


**Supplementary Figure 2. Detail of images registration.** (A) DAPI in each cycle is obtained. (B) Morphology transform is used to expose key points and to find matched key point pairs between images of cycle 1 and each other cycles. (C) Key point pairs are then filtered iteratively until no outlier was found. (D) At last the matched pairs will be used to calculate transform matrix in each registration, and matrix can be used in rigid transformation for images in their own cycles.

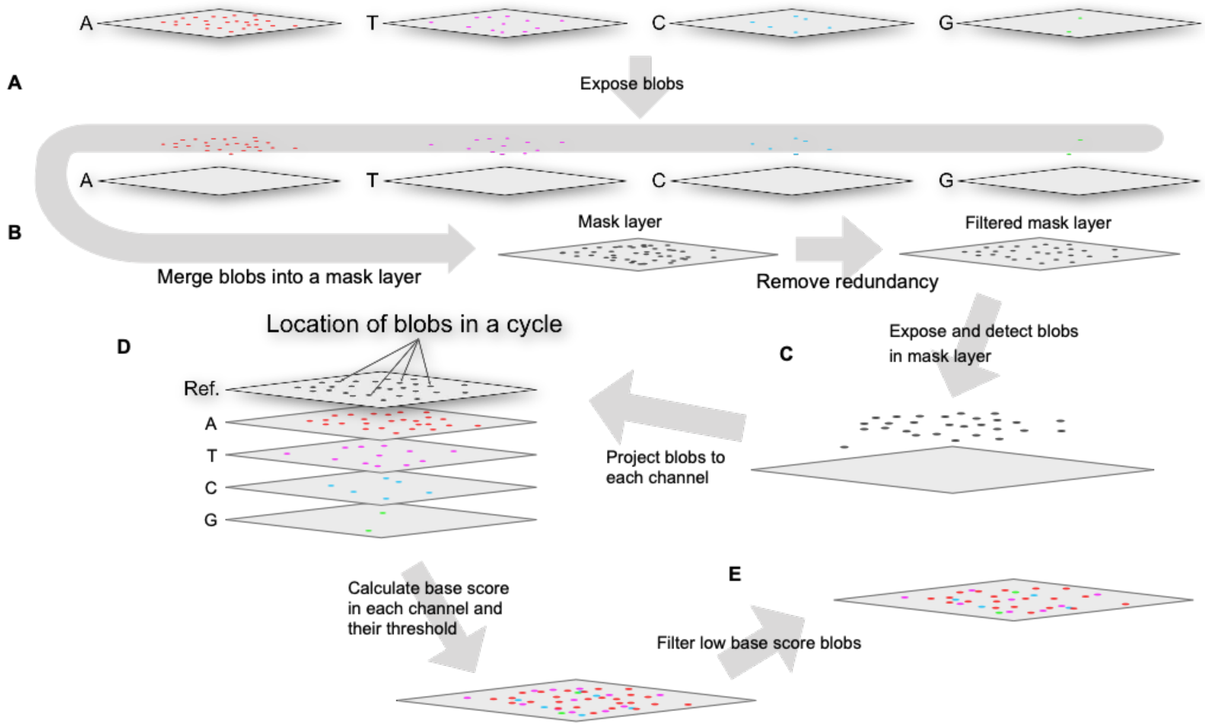


**Supplementary Figure 3. Detail of blob detection.** (A) Blobs are first exposed by morphological transform and detected. (B) Detected blobs in all channel from the same cycle are then projected into a mask layer. (C) Blobs are detected again on the mask layer. (D) Blobs' locations are projected to each channel and their base scores are calculated. Base score threshold for each channel in each cycle is taken as $round\left(\frac{mode_{base\_score}}{5}\right) * 5$. (E) Detected blobs are at last filtered based on the threshold.

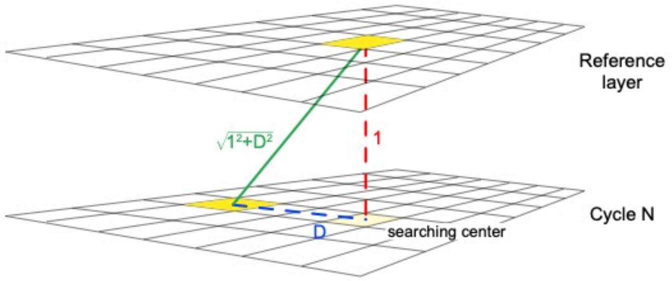


**Supplementary Figure 4. Base connection and error rate calibration.** Because the detected blobs' location might not be accurately aligned at pixel level, we projected the non-redundant blob from the reference cycle to every cycle, and search for candidate linked base at every cycle at the 6x6 region of the searching center. Error rate of the searched base at cycle N would be penalized by multiplying it by $\sqrt{1^2 + D^2}$, where the distance between reference layer at cycle N is defined to be one, and the distance between a searched candidate and searching center is defined to be D.

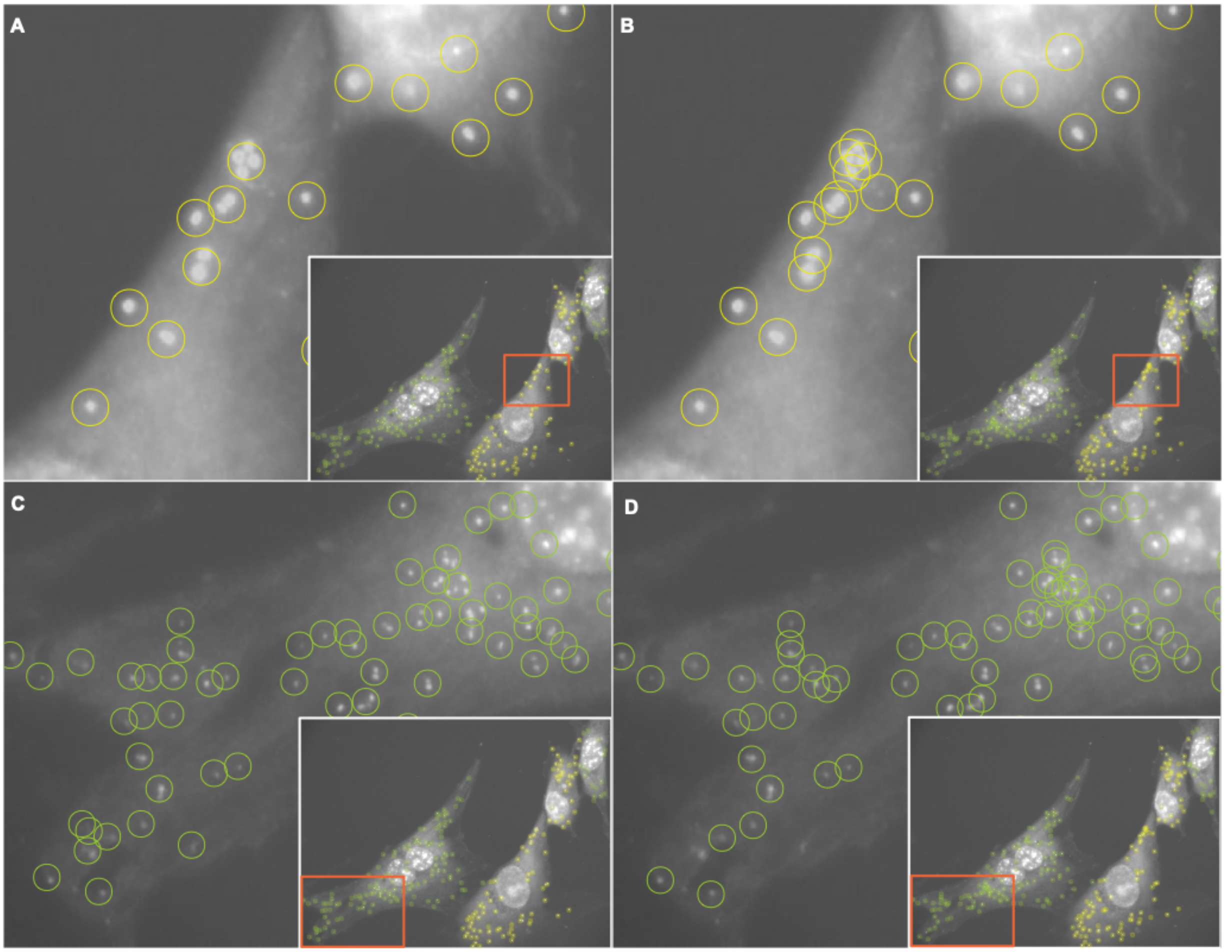


**Supplementary Figure 5. Comparison of detected barcode location in example regions of HM between IRIS and Ke's pipeline.** A, C and B, D show barcode location detected by Ke's CellProfiler pipeline and IRIS, respectively. Cycle 1 merged channels are used as background, while yellow and green circles indicate the detected location of two barcodes (yellow for barcode AGGC and green for barcode AAGC). Comparing A with B, a number of dense barcodes can be detected clearly by IRIS but Ke's pipeline confuse them as one. This same observation is also found in another region of this image shown in C and D. These cases show IRIS is sensitive to barcode-dense regions.

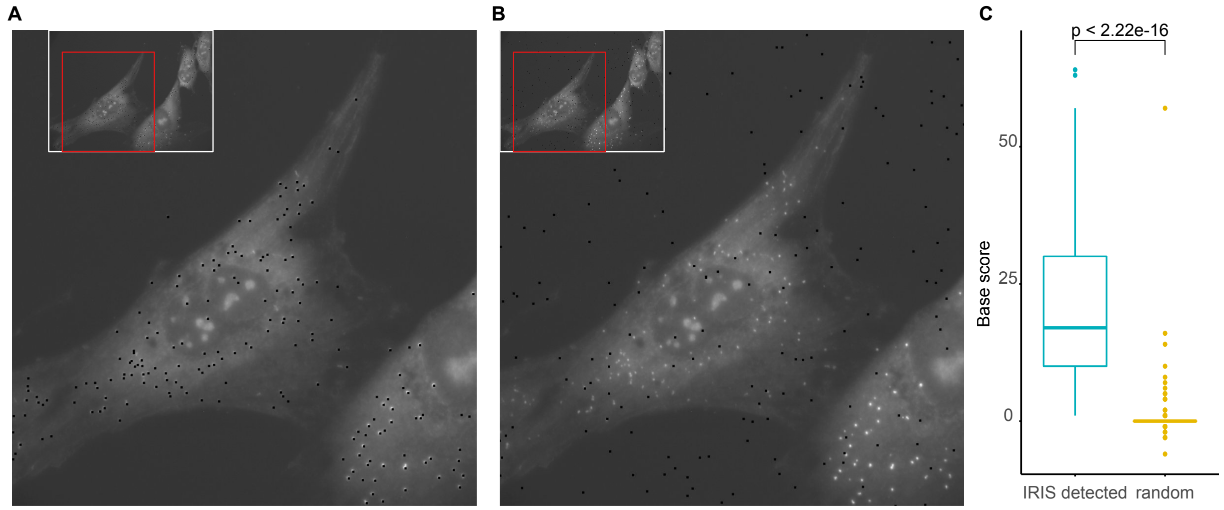


**Supplementary Figure 6. Detected barcodes spatial accuracy evaluation.** To evaluate location accuracy of IRIS, we compared the greyscale change between IRIS detected barcodes (A) and randomly selected blobs (B). Sampled locations are highlighted as black dots. (C) Spatial accuracy is qualified by comparing base score distribution between IRIS detected barcodes and random blobs. IRIS detected barcodes of HM data have significantly higher base score than random blobs, suggesting the accuracy of IRIS detected barcodes location.

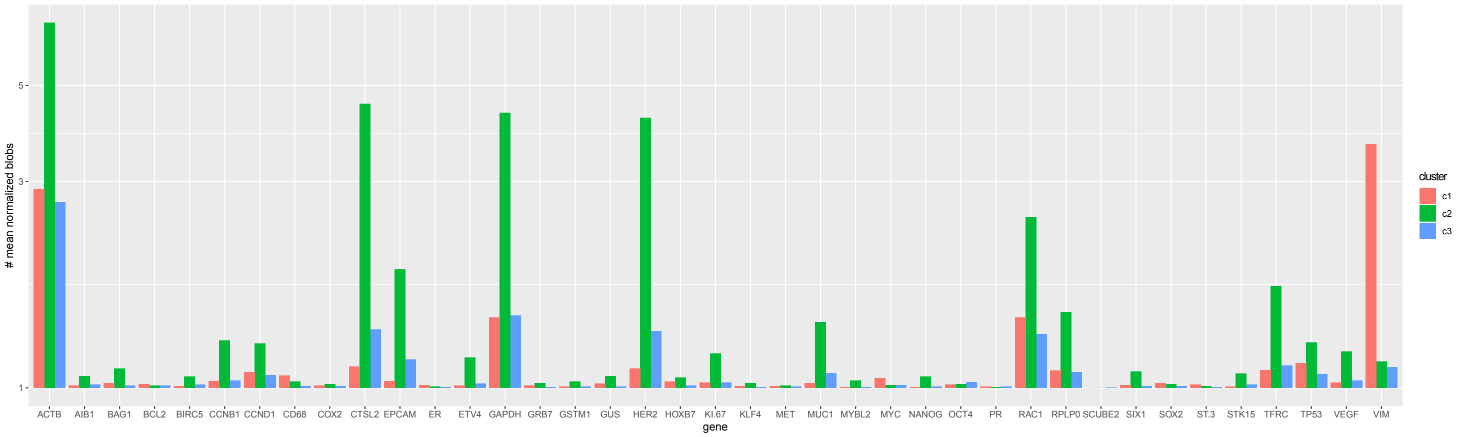


**Supplementary Figure 7. K-means cluster result in breast tumor slice data.** In this case, windows in cluster ‘c1’ and ‘c2’ are treated as ‘VIM+’ and ‘HER2+’ region in later correlation analysis, respectively.

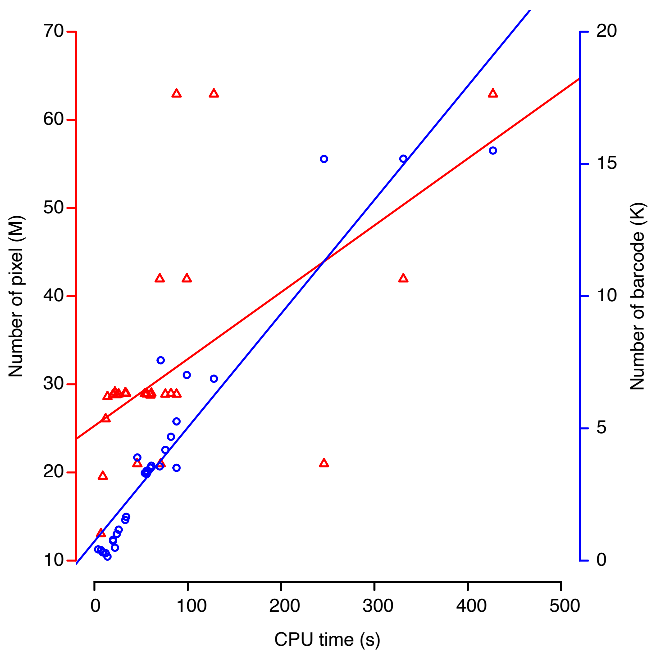


**Supplementary Figure 8. IRIS CPU time vs barcode number and image dataset size.** The fitted line between CPU time and the number of detected barcodes is steeper than that between CPU time and total number of pixels, suggesting that detected barcode number affect IRIS run time more than image size.

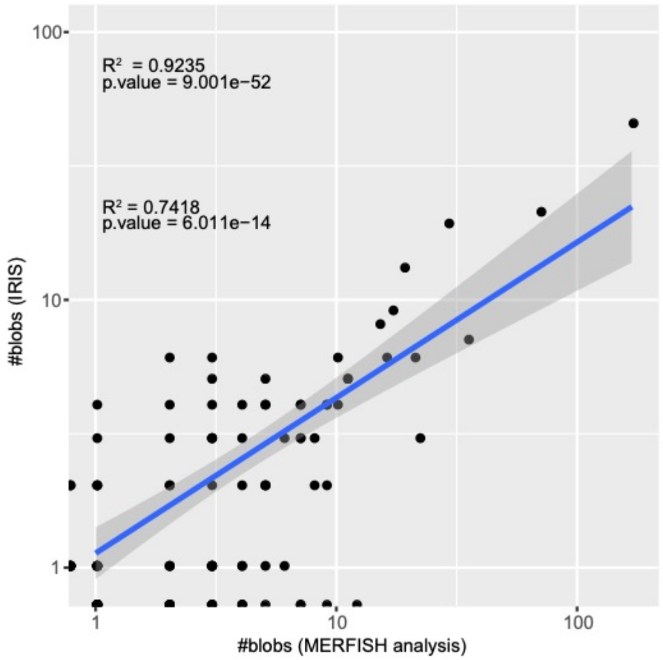


**Supplementary Figure 9. Correlation of detected barcode numbers between MERFISH analysis and IRIS.** The high correlation between these two datasets suggested IRIS' compatibility to MERFISH data. The $R^2$ and p-value above is calculated between raw barcode counts of each gene while the $R^2$ and p-value below is calculated between barcode counts after excluding zeros and log10 transformation.

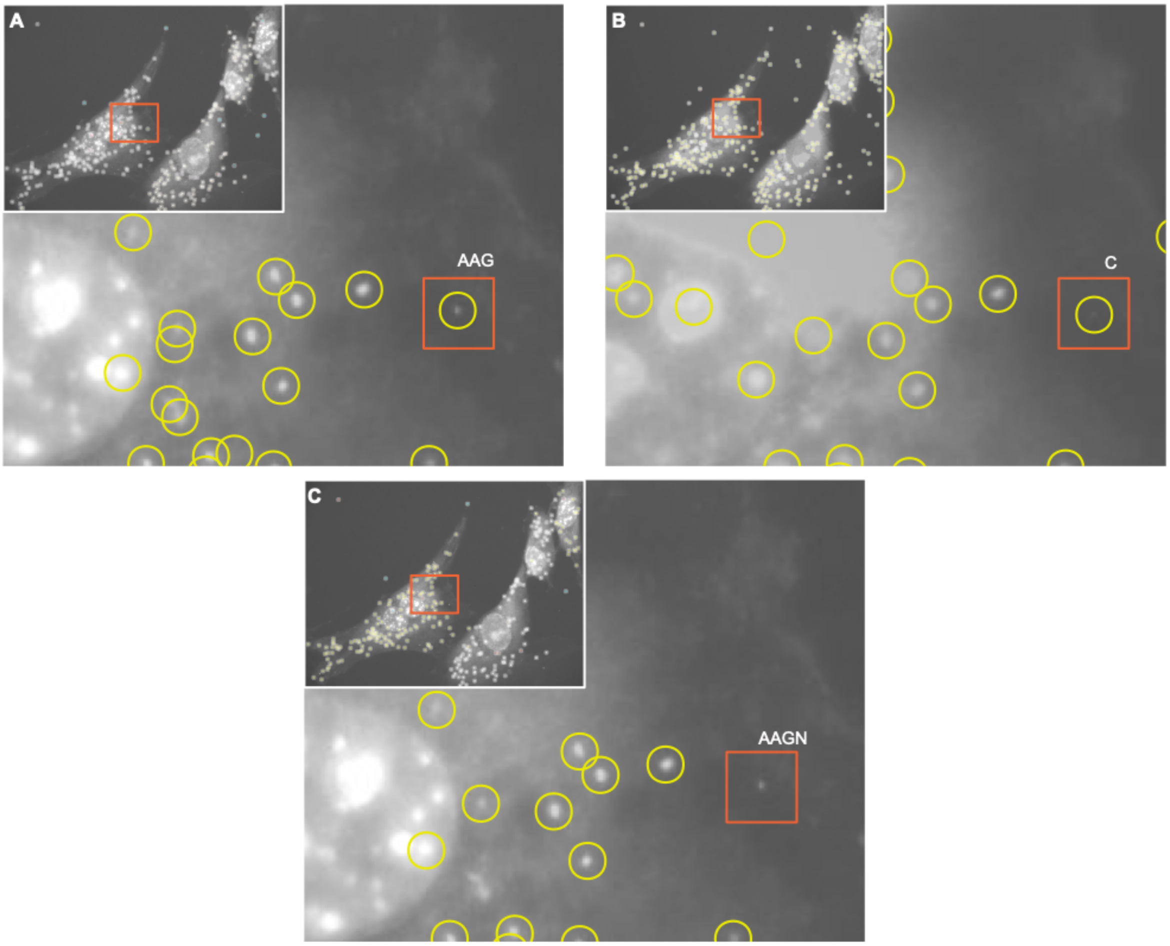


**Supplementary Figure 10. Imperfection of registration might cause base linking issue.** A minor change of blob position would cause barcode linking issue, thus the blob in the red frame could be detected and decoded containing no ‘N’ with images of cycle 1-3 (A) and images of cycle 4 only (B), but would contain an ‘N’ in the last base with all images from cycle 1-4 (C).

### Supplementary Tables

**Supplementary Table 1. Information of samples used in this study.**

| sample type | sample name | #slices | #view per slice | #cycle | resolution per image * | data availability |
|---|---|---|---|---|---|---|
| co-culture of human and mouse cell | HM | 1 | 1 | 4 | 1330x980 | http://d1zymp9ayga15t.cloudfront.net/content/Examplezips/ExampleInSituSequencing.zip |
| breast tumor slice | breast tumor slice | 1 | 16 | 4 | 1389x1041 | upon request |
| cell | MERFISH | 1 | 1 | 16 | 256x256 | http://zhuang.harvard.edu/MERFISHData/MERFISH_Examples.zip |

* images resolution in the same sample might vary a little

**Supplementary Table 2. Breast tumor slice data used for comparison among results of IRIS, Ke et al. 2013 and RNA-seq.**

| gene | VIM+ region | | normal breast TPM | HER2+ region | | BT474 cell line TPM |
|---|---|---|---|---|---|---|
| | #blobs (IRIS) | #blobs (Ke et al. 2013) | | #blobs (IRIS) | #blobs (Ke et al. 2013) | |
| ACTB | 699 | 1,053 | 2,826.19 | 2,278 | 3,335 | 1,908.75 |
| AIB1 | 4 | 5 | 5.56 | 25 | 4 | 146.70 |
| BCL2 | 7 | 18 | 50.17 | 5 | 2 | 17.42 |
| BIRC5 | 3 | 5 | 0.49 | 24 | 28 | 72.82 |
| CCNB1 | 14 | 14 | 2.83 | 109 | 90 | 40.81 |
| CCND1 | 32 | 18 | 408.09 | 101 | 86 | 394.01 |
| CD68 | 25 | 25 | 225.33 | 13 | 4 | 3.77 |
| EPCAM | 14 | 35 | 4.53 | 334 | 546 | 71.88 |
| ETV4 | 4 | 3 | 4.04 | 67 | 69 | 6.98 |
| GAPDH | 168 | 281 | 1,189.59 | 1,266 | 1,682 | 1,666.38 |
| GUS | 8 | 1 | 53.62 | 25 | 6 | 51.03 |
| HER2 | 40 | 60 | 35.03 | 1,223 | 1,948 | 2511.91 |

| | | | | | | |
|---|---|---|---|---|---|---|
| MET | 3 | 2 | 3.88 | 5 | 3 | 5.35 |
| MUC1 | 9 | 6 | 8.98 | 160 | 138 | 13.75 |
| MYBL2 | 1 | 0 | 2.80 | 15 | 0 | 102.99 |
| MYC | 20 | 7 | 145.10 | 6 | 2 | 158.50 |
| RAC1 | 168 | 202 | 312.72 | 564 | 669 | 433.25 |
| RPLP0 | 36 | 33 | 266.30 | 189 | 172 | 349.82 |
| SCUBE2 | 0 | 3 | 22.62 | 0 | 0 | 1.18 |
| SIX1 | 5 | 2 | 19.53 | 34 | 19 | 26.55 |
| STK15 | 2 | 4 | 0.00 | 30 | 37 | 121.41 |
| ST-3 | 0 | 6 | 2.01 | 0 | 1 | 4.67 |
| TFRC | 37 | 50 | 7.19 | 275 | 343 | 86.08 |
| TP53 | 52 | 69 | 25.46 | 104 | 150 | 81.22 |
| VEGF | 11 | 24 | 27.07 | 81 | 143 | 31.82 |
| VIM | 987 | 901 | 2,399.79 | 57 | 23 | 46.18 |

**Supplementary Table 3. IRIS' computational performance on available ISS data.** 'size of DAPI in cycle1' means image resolution of cycle 1 DAPI, other image in this run should be similar with it; '#cycles' means the number of cycles we used in a single run; 'total pixels' is the sum of pixels of all images in a single run in our test; '#called barcodes (no N)' means the number of IRIS detected barcodes containing no "N".

| sample name | size of DAPI in cycle1 | #cycles | total pixels | #called barcode (no N) | CPU TIME (s) |
|---|---|---|---|---|---|
| HM | 1330 x 980 | 1 | 6,517,000 | 424 | 3.780 |
| HM | 1330 x 980 | 2 | 13,034,000 | 397 | 7.408 |
| HM | 1330 x 980 | 3 | 19,551,000 | 311 | 9.326 |
| HM | 1330 x 980 | 4 | 26,068,000 | 282 | 11.663 |
| breast tumor slice (slideA_1) | 1389×1041 | 4 | 28,761,335 | 3,514 | 59.886 |
| breast tumor slice (slideA_2) | 1390×1042 | 4 | 28,870,440 | 5,263 | 87.546 |
| breast tumor slice (slideA_3) | 1391×1043 | 4 | 28,943,390 | 4,682 | 81.997 |
| breast tumor slice (slideA_4) | 1392×1044 | 4 | 29,004,165 | 1,536 | 33.393 |
| breast tumor slice (slideA_5) | 1395×1047 | 4 | 29,101,705 | 487 | 22.374 |
| breast tumor slice (slideA_6) | 1392×1044 | 4 | 28,979,805 | 3,588 | 60.608 |
| breast tumor slice (slideA_7) | 1388×1040 | 4 | 28,931,165 | 3,313 | 53.920 |

| | | | | | |
|---|---|---|---|---|---|
| breast tumor slice (slideA_8) | 1391×1043 | 4 | 28,882,645 | 3,394 | 55.511 |
| breast tumor slice (slideA_9) | 1390×1042 | 4 | 28,943,300 | 3,273 | 55.848 |
| breast tumor slice (slideA_10) | 1390×1042 | 4 | 28,967,690 | 1,655 | 33.883 |
| breast tumor slice (slideA_11) | 1390×1042 | 4 | 28,834,215 | 745 | 19.736 |
| breast tumor slice (slideA_12) | 1392×1044 | 4 | 28,810,035 | 788 | 20.310 |
| breast tumor slice (slideA_13) | 1384×1036 | 4 | 28,797,650 | 1,005 | 24.068 |
| breast tumor slice (slideA_14) | 1386×1038 | 4 | 28,591,945 | 151 | 13.957 |
| breast tumor slice (slideA_15) | 1390×1042 | 4 | 28,882,605 | 1,170 | 26.310 |
| breast tumor slice (slideA_16) | 1390×1042 | 4 | 28,870,440 | 4,186 | 75.851 |

* Platform: MacBookPro11,4 (Mid 2015); CPU: 2.2 GHz Quad-Core Intel Core i7; Mem: 16 GB 1600 MHz DDR3

**Supplementary Table 4. Blobs number of MERFISH example data called by MERFISH analysis and IRIS.**

| gene | #blobs (MERFISH_analysis) | #blobs (IRIS) |
|---|---|---|
| ABCA2 | 1 | 1 |
| AFAP1 | 9 | 4 |
| AFF4 | 7 | 0 |
| AGPS | 1 | 1 |
| AKAP11 | 2 | 0 |
| ALMS1 | 2 | 0 |
| ALPK2 | 1 | 0 |
| AMOTL1 | 6 | 1 |
| ANKRD52 | 7 | 4 |
| ATP11A | 0 | 1 |
| C17orf51 | 1 | 1 |
| CASC5 | 1 | 0 |
| CBX5 | 3 | 1 |
| CENPF | 5 | 1 |
| CEP250 | 2 | 1 |
| CHD8 | 2 | 0 |
| CKAP5 | 7 | 4 |
| COL5A1 | 11 | 5 |
| COL7A1 | 4 | 0 |
| CREBBP | 4 | 2 |
| DIEXF | 3 | 3 |
| DIP2B | 1 | 1 |
| DIP2C | 5 | 2 |
| DNAJC13 | 5 | 0 |
| DOCK7 | 1 | 1 |
| DOPEY1 | 1 | 0 |
| DSEL | 1 | 0 |
| DYNC1H1 | 16 | 6 |
| EGFR | 3 | 2 |
| FAM208B | 4 | 4 |

| | | |
|---|---|---|
| FASN | 1 | 0 |
| FBN1 | 7 | 3 |
| FBN2 | 10 | 4 |
| FLNA | 169 | 45 |
| FLNB | 2 | 0 |
| FLNC | 22 | 3 |
| FRY | 0 | 1 |
| FYCO1 | 3 | 0 |
| FZD4 | 1 | 0 |
| GPR107 | 4 | 0 |
| GTF3C1 | 2 | 0 |
| GTF3C4 | 0 | 1 |
| H6PD | 5 | 2 |
| HEATR5B | 1 | 2 |
| HELZ2 | 0 | 1 |
| HERC2 | 4 | 1 |
| HIVEP1 | 1 | 3 |
| IGF2R | 8 | 2 |
| IQGAP1 | 10 | 6 |
| ITGA2 | 1 | 1 |
| KIAA1147 | 1 | 0 |
| KIAA1199 | 5 | 5 |
| KIAA1462 | 7 | 3 |
| KIAA2018 | 0 | 1 |
| KIDINS220 | 0 | 1 |
| KPNA4 | 2 | 0 |
| LMTK2 | 2 | 1 |
| LRP1 | 9 | 4 |
| LUZP1 | 4 | 0 |
| LYST | 0 | 1 |
| MALAT1 | 1 | 0 |
| MAN1A2 | 1 | 0 |
| MED14 | 0 | 2 |
| MKI67 | 17 | 9 |
| MYH10 | 15 | 8 |
| NF1 | 3 | 0 |
| NOTCH2 | 2 | 2 |
| NRIP1 | 2 | 1 |
| NUMA1 | 0 | 1 |
| NUP98 | 5 | 4 |
| PAPPA | 2 | 2 |
| PCNX | 3 | 3 |
| PDS5A | 2 | 3 |
| PHIP | 1 | 1 |
| PIEZO1 | 4 | 1 |
| PLXNA1 | 3 | 3 |
| POLQ | 1 | 2 |
| PRKCA | 11 | 5 |
| PRKDC | 3 | 6 |
| PRPF8 | 29 | 19 |
| PRRC2B | 1 | 2 |
| PTPN14 | 7 | 0 |
| RAB3B | 9 | 0 |
| RNF169 | 1 | 4 |
| ROCK2 | 3 | 1 |
| SCUBE3 | 1 | 0 |
| SIPA1L3 | 5 | 4 |
| SLC5A3 | 1 | 0 |
| SLC7A11 | 1 | 0 |
| SLC38A1 | 4 | 3 |
| SMARCA5 | 2 | 6 |

| | | |
|---|---|---|
| SON | 5 | 2 |
| SPTAN1 | 21 | 6 |
| SPTBN1 | 3 | 1 |
| SRRM2 | 19 | 13 |
| SSH1 | 3 | 1 |
| STARD9 | 0 | 2 |
| SVEP1 | 1 | 0 |
| TEAD1 | 9 | 2 |
| THBS1 | 70 | 21 |
| THSD4 | 3 | 1 |
| TLN1 | 35 | 7 |
| TMOD2 | 1 | 1 |
| TNC | 12 | 0 |
| TNRC6A | 3 | 5 |
| TPR | 8 | 3 |
| UBR2 | 3 | 4 |
| UBR5 | 3 | 1 |
| USP9X | 1 | 0 |
| USP24 | 3 | 0 |
| USP34 | 6 | 3 |
| VCAN | 3 | 1 |
| VPS13A | 0 | 1 |
| VPS13D | 3 | 0 |
| ZC3H13 | 2 | 0 |
| ZFC3H1 | 1 | 1 |
| ZKSCAN2 | 2 | 4 |
| blank001 | 1 | 0 |
| notarget001 | 1 | 0 |
| notarget004 | 0 | 1 |
| notarget005 | 0 | 2 |
